# SomaMutDB 2.0: A comprehensive database for exploring somatic mutations and their functional impact in normal human tissues

**DOI:** 10.1101/2025.09.14.676124

**Authors:** Anthony Shea, Shixiang Sun, Justin Kennedy, Lei Zhang, Jan Vijg, Xiao Dong

## Abstract

Recent advances in ultra-accurate sequencing technologies have revealed that somatic mutations accumulate across the human lifespan and may contribute to both normal aging and disease. These mutations are highly diverse, often non-recurrent, and functionally heterogeneous, which makes their biological impact difficult to evaluate systematically. Although many studies have profiled somatic mutations in individual tissues or limited cohorts, a centralized and scalable platform that integrates discoveries and supports functional interpretation has been lacking. To address this gap, we present SomaMutDB 2.0 (https://somamutdb.org/SomaMutDB/), a substantially expanded database that catalogs 8.9 million mutations (8.57 million SNVs and 0.29 million INDELs) from 10,852 samples of 607 human subjects across 47 studies. Beyond expanded data coverage, SomaMutDB 2.0 introduces a comprehensive functional annotation framework that applies 22 predictive models, encompassing coding, regulatory, expression-based, and ensemble predictors, to systematically assess mutational impact. Users can browse pre-annotated variants through an interactive interface or upload their own variants for real-time analysis, with results contextualized against all mutations from normal, non-diseased tissues in the database. Together, these advances establish SomaMutDB 2.0 as the most comprehensive resource currently available for characterizing somatic mosaicism and functional interpretation in human health and aging.

**Graphical Abstract:** SomaMutDB 2.0 provides an expanded catalog of 8.9 million somatic mutations across 30 tissues, along with pipelines for mutational signature analysis and 22-tool functional annotation that enable user-submitted variant interpretation.

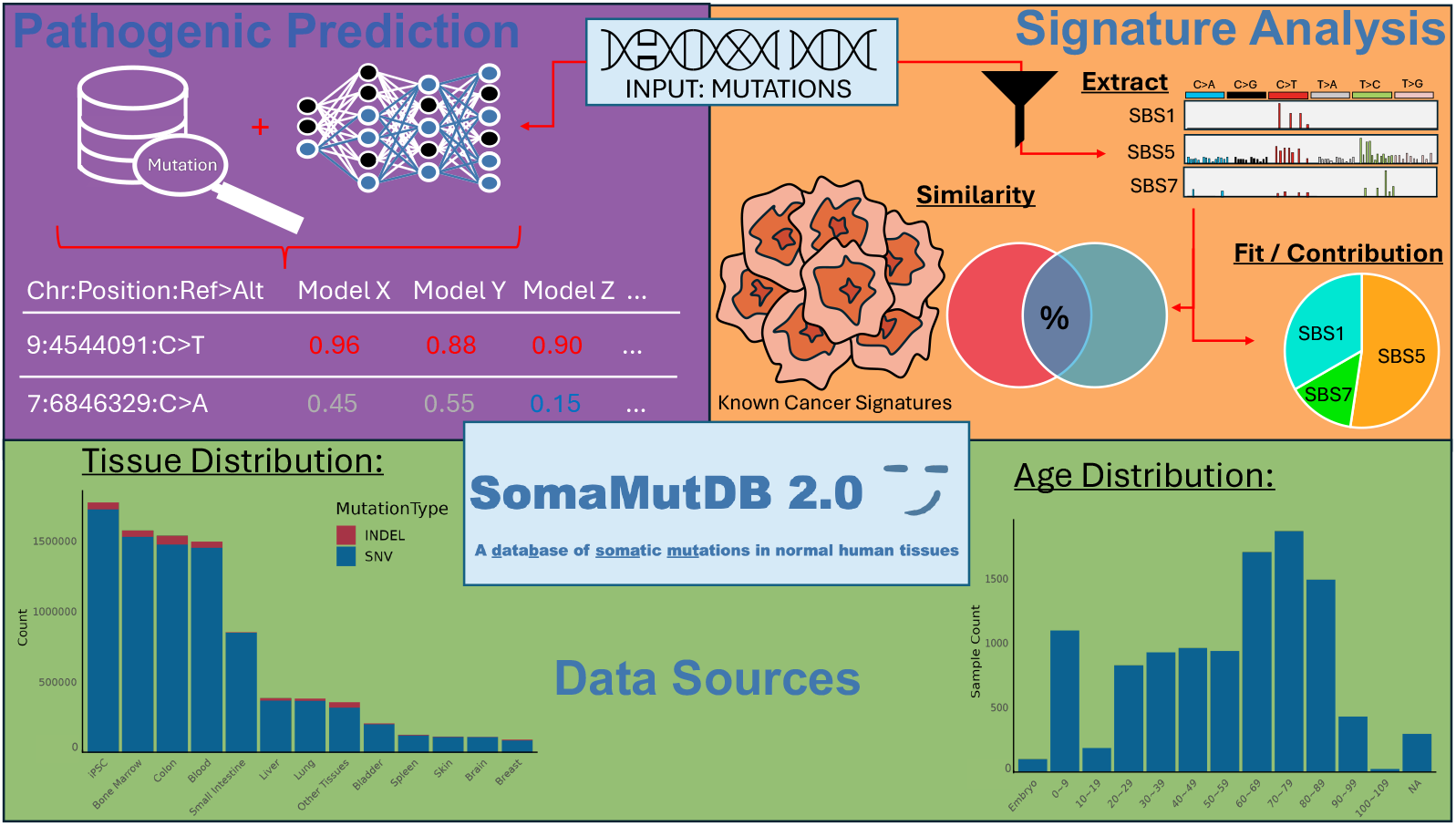

## Introduction

Somatic mutations are well-established drivers of cancer, but their roles in other chronic diseases and in age-related cellular decline in normal human tissues remain difficult to study. Because the error rate of standard DNA sequencing is several orders of magnitude higher than the frequency of somatic mutations, only germline variants and somatic mutations that have undergone clonal expansion can be reliably detected by bulk sequencing. Consequently, most large-scale analyses to date have focused on tumors, where clonal expansion enables accurate cataloging of mutation frequencies and signatures, as exemplified by The Cancer Genome Atlas (TCGA) (1), the International Cancer Genome Consortium (ICGC) (2), and the Catalogue of Somatic Mutations in Cancer (COSMIC) (3).

To overcome the challenge of analyzing somatic mutations in noncancerous tissues, more accurate sequencing approaches have been developed, including single-cell DNA sequencing (4,5), deep sequencing of natural or *in vitro* clones (6,7), and single-molecule (duplex) sequencing (8,9). These technologies have enabled recent discoveries of age- and chronic disease-associated somatic mutation accumulation across multiple human tissues (10-12). However, because of their high cost, individual studies typically focused on a single tissue type and analyzed samples from only 10–20 subjects. This limitation motivated us to develop SomaMutDB v1.0, a catalog comprising 2.42 million somatic single-nucleotide variants (sSNVs) and 0.12 million somatic small insertions and deletions (sINDELs) identified in 19 normal tissues and cell types, based on 2,838 single cells, clones, or biopsies from 374 human subjects (13).

A major challenge in characterizing somatic mutagenesis in normal cells is that, unlike in tumors, the same mutations are rarely observed recurrently across different samples or studies. This is not unexpected, as most single-cell level mutations arise from stochastic errors in DNA damage or replication. These events appear largely random, yet they still display characteristic patterns such as mutational spectra and signatures (11,14). Although it may be impossible to experimentally validate the functional impact of every mutation and their interactions, substantial progress in computational annotation, particularly through deep learning–based approaches, has enabled large-scale evaluation of mutational consequences (15). At the same time, research on human somatic mosaicism has expanded rapidly. Large-scale consortium initiatives, such as the Somatic Mosaicism across Human Tissues (SMaHT) Network (16), together with population-wide epidemiological studies (17) powered by single-molecule sequencing technologies, are now uncovering somatic mutations in normal tissues at an unprecedented scale.

Inspired by these developments, we present SomaMutDB 2.0, which substantially expands both the scope and utility of the database. The updated catalog now contains 8.9 million mutations (8.57 million sSNVs and 0.29 million sINDELs) from 47 studies, i.e., a 3.5-fold increase over version 1.0. This expansion reflects the accelerating pace of discovery enabled by emerging technologies such as Nanorate sequencing (NanoSeq) (9) and Single-Molecule Mutation Sequencing (SMM-seq) (18). Beyond data growth, SomaMutDB 2.0 introduces a comprehensive functional annotation framework that integrates 22 computational prediction tools, ranging from widely used algorithms (e.g., SIFT4G (19), CADD (20)) to state-of-the-art deep learning methods (e.g., AlphaMissense (21), EVE (22), PrimateAI (23)). In addition, users can upload their own variants for on-demand functional assessment and mutational signature analysis. This system supports systematic evaluation of both coding and regulatory variants by providing absolute impact scores as well as contextualized information through relative rankings and score distributions across all SomaMutDB mutations, offering a more interpretable view of mutational consequences. Together, these advances establish SomaMutDB 2.0 as the most comprehensive and versatile resource for characterizing somatic mutations in normal human tissues, effectively bridging the gap between mutation discovery and functional interpretation.

## Materials and Methods

### System design and implementation

SomaMutDB 2.0 was built on the same database and web frameworks as version 1.0 (13). For the new user-submission function of mutation annotation, queries are processed through a job scheduler (Sun Grid Engine, SGE; version 8.1.9). Once the analysis is completed, SomaMutDB 2.0 automatically sends the user an email containing a query ID, which can be used to retrieve the results.

### Data sources

Since the release of SomaMutDB 1.0, the database has been expanded by incorporating somatic mutations from 23 additional studies published in the past three years. Consistent with the criteria used in version 1.0, we included sSNVs and sINDELs data derived from various sample types, e.g., single cell, clone, and tissue biopsy, with either whole genome or whole exome coverage. From our search and visualization functions, we excluded samples from tumors and other abnormal tissues, retaining only somatic mutations identified in normal tissues and cell types, although the mutations from tumors and other abnormal tissues are available for download from the database. Examples of excluded samples include those from Alzheimer’s disease (9,24), alcoholic liver cirrhosis (25), and antiviral treatment in a transplantation recipient (26). In addition to the discovery strategies incorporated previously, the new version also integrates mutations identified using duplex sequencing protocols, i.e., NanoSeq (9) and SMM-seq (18).

### Data processing

For the newly added studies, all reported mutations were processed using the standard pipeline established in SomaMutDB 1.0, with exceptions in three specific scenarios. First, for studies in which mutations were present in different format, we now changed them to ensure consistency, e.g., CTT>CT at chrA:positionB is now shown as TT>T at chrA:positionB+1, or CTAGTC>CGATC is now shown as CTAG>CGA in SomaMutDB (25,27). Second, double-nucleotide variants were decomposed into individual single-nucleotide variants for consistency (28,29). Third, one study did not provide subject-level information; in this case, variants are presented as they were listed in the supplementary table of the original paper (30). Mutations located in small contigs of the reference genome (not assigned to specific chromosome positions) were excluded, while only those mapped to chromosomes 1–22, X, and Y were retained.

### Previous features in 2.0

All features from the 1.0 version of the database have been updated to incorporate the new functional annotations. In addition, the legacy version of the website remains accessible to all users at its original domain (https://vijglab.einsteinmed.org/SomaMutDB).

### A comprehensive platform to annotate mutational impact

To provide comprehensive functional annotation, SomaMutDB 2.0 incorporates a computational pipeline that runs 22 established prediction models (**Table S1**). We chose these models because they can cover broadly four categories based on their predictive scope. The first are Protein Impact Models (PIMs), which evaluate the functional consequences of coding variants by leveraging evolutionary conservation, amino acid properties, and structural features to classify mutations as benign or deleterious. The second category comprises Regulatory and Non-coding Impact Models (RNIMs), which assess the potential effects of variants located in promoters, enhancers, splice sites, and other regulatory regions that may alter gene expression. The third group is a Gene Expression Predictor (GEP; only one is available in this group, i.e., ExPecto (31)), which directly estimates how variants influence the expression of their nearest genes in various tissue types. Finally, Meta-Predictors (MPs) integrate outputs from multiple algorithms in an ensemble framework to generate consensus scores that improve predictive accuracy by leveraging the strengths of individual methods.

Because many of these tools are integrated into the Ensembl Variant Effect Predictor (VEP) (32), we implemented VEP (version 113) with the dbNSFP plugin (version 4.7) (33) locally on the SomaMutDB server (using GRCh38 coordinates). The dbNSFP plugin provides access to 17 models spanning the categories described above, including AlphaMissense (21), CADD (20), DANN (34), DEOGEN2 (35), EVE (22), MPC (36), and PrimateAI (23) for PIMs; Eigen (37), GenoCanyon (38), and LIST-S2 (39) for RNIMs; and BayesDel (40), M-CAP (41), MetaRNN (42), MetaSVM (43), MetaLR (43), MVP (44), and REVEL (45) as ensemble predictors MPs. In addition to raw outputs, dbNSFP also reports rank scores derived from its internal mutation database, enabling variants to be compared within a broader genomic context.

Five further tools were implemented individually to extend coverage beyond the dbNSFP framework. FATHMM provides conservation-based pathogenicity predictions with pre-computed hg19 scores for both coding and non-coding variants (46). CanDrA is a cancer-specific pathogenicity model run with the pan-cancer GENERAL dataset under hg19 coordinates (47). FINSURF annotates non-coding variants and produces both a numeric score and a feature-contribution plot integrating sequence, conservation, epigenomic, and gene association data (48). MutationTaster2025, accessed via the GeneCascade API, combines conservation and functional annotation features, with transcript-level consensus calling applied to classify variants as pathogenic, benign, or uncertain depending on whether ≥75% of transcripts support a given category (49). Finally, ExPecto is a tissue-specific model that estimates variant impact on gene expression across 200 tissues and cell types using hg19 coordinates (31).

All mutations in SomaMutDB have been annotated using this full set of 22 models. Users can also submit custom variant lists, which are processed automatically through the complete annotation framework. Jobs are managed with the Sun Grid Engine (SGE) scheduler, and UCSC LiftOver tool (hg19ToHg38.over.chain and hg38ToHg19.over.chain) ensures bidirectional compatibility across reference assemblies, with unmapped coordinates tracked and returned to users. Processing time scales linearly with variant count, with analysis of 1,000 variants typically completing within 20–30 minutes. Upon job completion, users receive an automated email with instructions to access their results. Importantly, SomaMutDB provides not only the raw outputs of each prediction model but also contextualized information for every mutation, including its relative ranking and score distribution across all variants in the SomaMutDB database, thereby offering a more interpretable view of mutational consequences.

## Results

### Database content

SomaMutDB continues to serve as a dedicated database and analytical platform for somatic mutations identified in normal human tissues and cells. In version 2.0, the database contains 8,866,453 mutations (8,576,771 SNVs and 289,682 INDELs) derived from 10,852 samples, including single cells, clones, and biopsies, collected from 607 human subjects across 47 studies. Because age remains the most significant factor influencing mutation burden and has been the focus of many recent studies, the samples included in SomaMutDB 2.0 cover a broad and balanced distribution of age groups, ranging from embryonic stages to centenarians (**Figure 1A**). The cohort is approximately 46% female and 53% male (**Figure 1B**), and spans 30 tissue and cell types, another key determinant of mutational burden, spectrum, and clonality (**Figure 1C; Table S2**).

**Figure 1.**
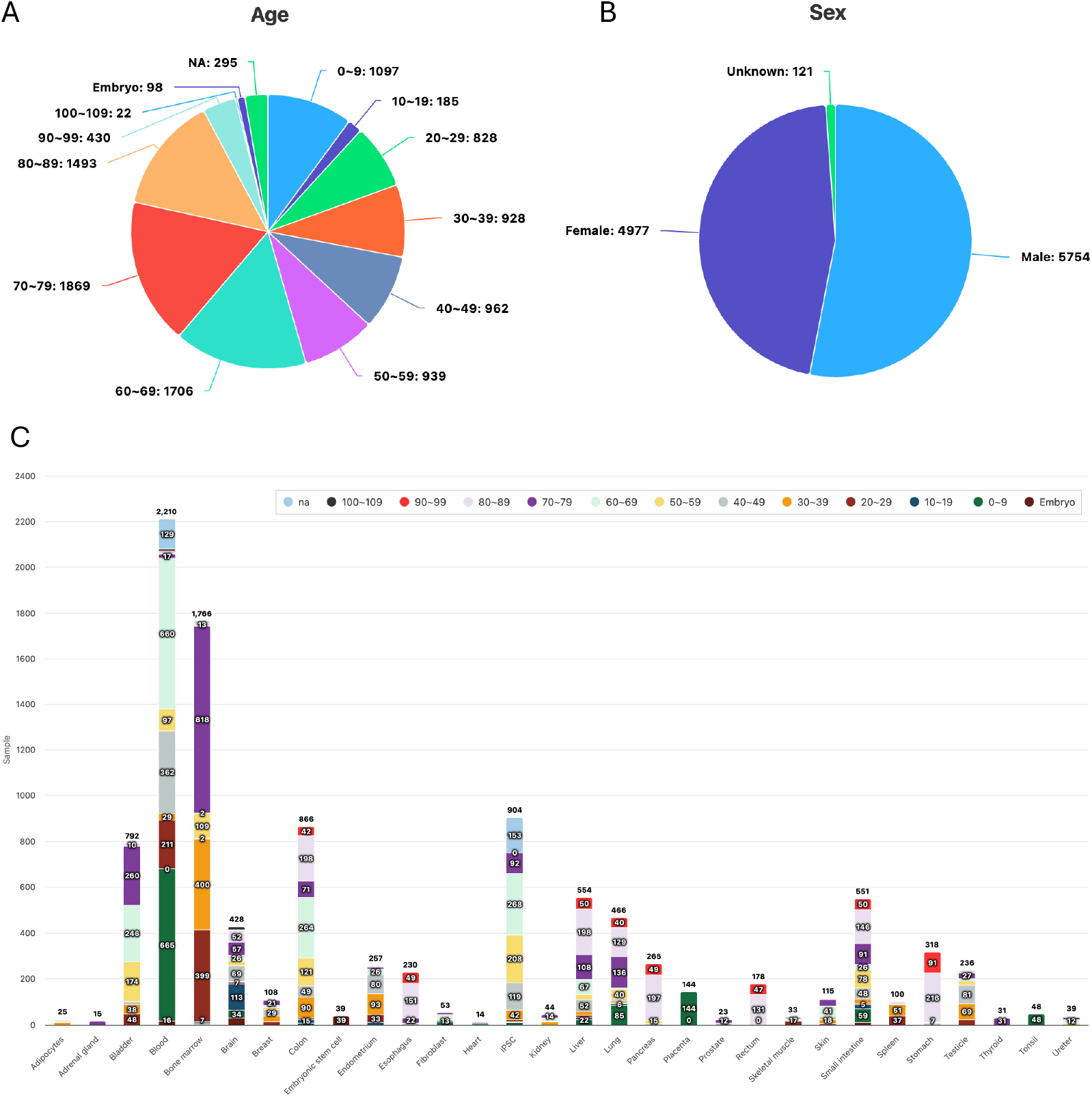
Distribution of mutations by age, sex, and tissue type. (A) Pie chart showing mutation distribution across age groups from embryonic samples to centenarians. (B) Pie chart showing mutation distribution by sex. (C) Bar plot showing mutation distribution across tissue types, with each bar subdivided by age group.

### Mutational impact prediction

In addition to the features available in version 1.0, such as the Genome Browser, Search functions, and Mutational Signature Analysis (13), the most significant advance in SomaMutDB 2.0 is the introduction of a comprehensive annotation framework for evaluating mutational impact using 22 complementary prediction algorithms of four categories (**Table S1**; **Materials and Methods**). This framework addresses the fact that no single algorithm can fully capture all mechanisms of pathogenicity, particularly for non-coding variants, whose effects are often context-dependent, mechanistically diverse, and potentially distal from the target gene.

Annotations for each mutation in SomaMutDB can be accessed in two ways. First, through the Genome Browser page, users can locate a mutation of interest (**Figure 2A**) and view its annotations directly in the “Attributes” section of the feature details panel (**Figure 2B**). Second, through the Search page, users can click the settings icon (gear symbol) to select which annotation algorithms to load (**Figure S1**), then query mutations based on gene, gene lists, genomic regions, or tissue type (**Figure 2C**). The search results display mutations in a tabular format, accompanied by their annotations, which can be directly downloaded on the same webpage (Download button).

**Figure 2.**
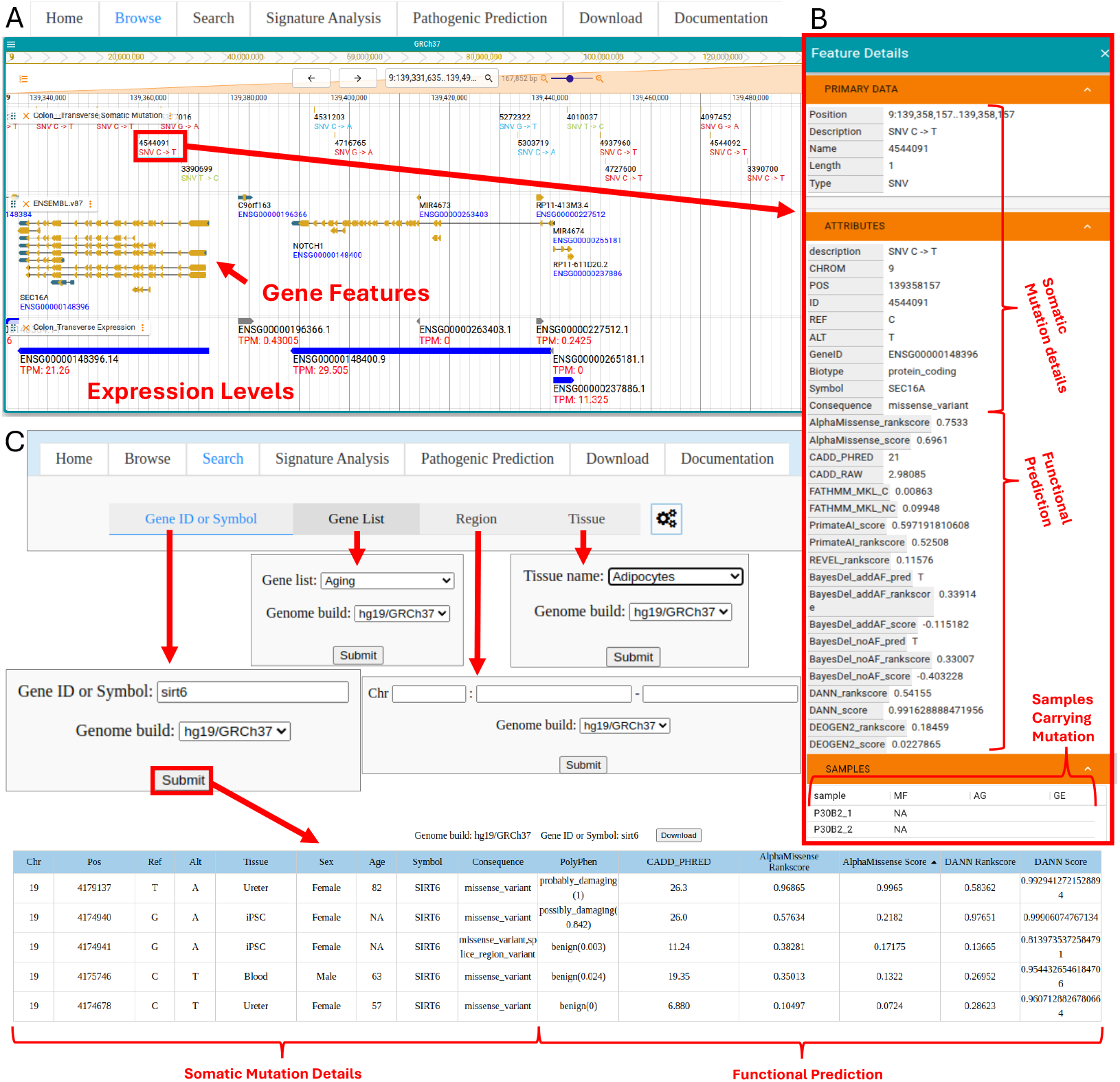
Updated browsing and search functionality. (A) Genome browser interface displaying gene annotations and expression tracks. Selection of a mutation triggers a feature details panel that reports mutation-specific attributes alongside functional predictions derived from integrated pathogenicity models. (B) Representative outputs from the search interface, shown for an example query of the SIRT6 locus. Search results return all mutations meeting the specified criteria, including variant descriptions, associated sample metadata, and selected functional predictions.

Beyond catalogued mutations, SomaMutDB 2.0 also allows users to upload their own variants for functional assessment. Submitted variants are processed through the full annotation pipeline, and the results are contextualized against the distribution of all mutations in SomaMutDB (derived from normal, non-diseased tissues), thereby enabling more informative interpretation of potential pathogenicity. This function is available through the new Pathogenetic Prediction page, where users can submit variants via file upload or text input, specify genome build, and provide a user’s email address (**Figure 3A**). Once analysis is complete, users are notified by both browser notification, which takes time to load, and/or email, which includes links to the results (**Figure 3B**).

**Figure 3.**
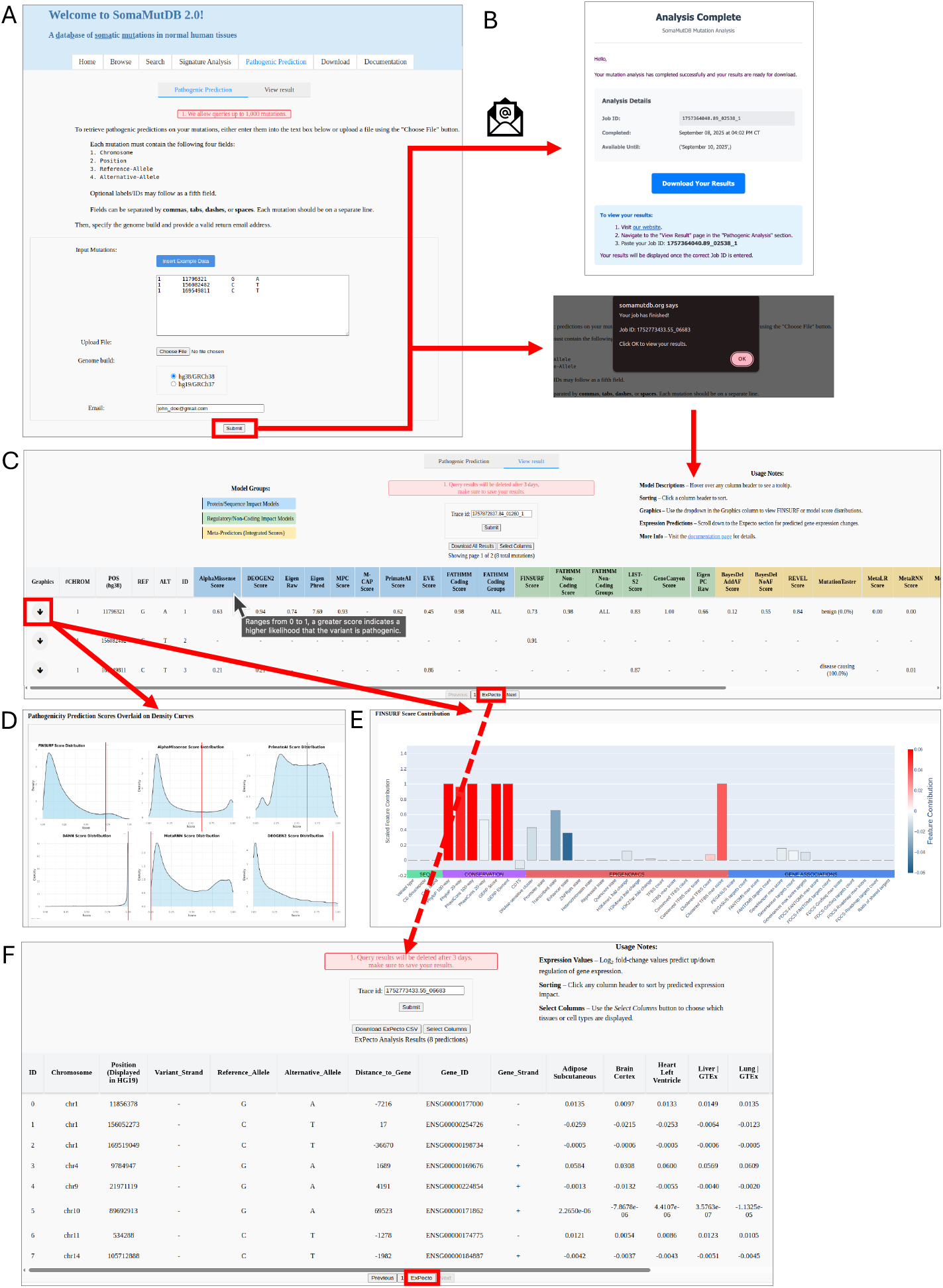
Pathogenic prediction query workflow and results interface. (A) Query input interface allowing users to enter mutations via text box or file upload, with required fields for genome build and user email. (B) An email and pop-up notification appear upon query completion. (C) Results retrieval using the returned trace ID displays all mutation functional predictions in a table organized by model type. (D) A graphics dropdown section shows score distributions retrieved from the SomaMutDB mutation library with the queried mutation score overlaid. (E) The graphics section also displays FINSURF feature contributions for mutations receiving FINSURF model scores. (F) ExPecto results accessible via tab navigation show log2 fold-change values for each mutation across selected tissue types.

Results are displayed in a table in which each column corresponds to a prediction model that generated a score (**Figure 3C**). Hovering over a column header reveals a tooltip with a brief description of the respective model (**Figure 3C**). Users can customize the output by choosing to display raw scores, ranked scores (derived from the dbNSFP database (33) rather than SomaMutDB), or both, through the “Select Columns” option (**Figure S2**). Columns are grouped and color-coded by model type, i.e., Protein Impact Models (PIMs), Regulatory and Non-Coding Impact Models (RNIMs), and Meta-Predictors (MPs). Dropdown arrows provide additional context, such as density plots showing the distribution of scores across all SomaMutDB mutations (**Figure 3D**). For one of the models, FINSURF with multiple feature-level outputs (48), additional details including contributions from sequence conservation, epigenomic marks, and nearest-gene associations, are available also through expandable rows (**Figure 3E**). Predictions from ExPecto (31), the only Gene Expression Predictor (GEP) in SomaMutDB, are displayed separately in the navigation bar below the main table to prevent overcrowding, as the results include multiple columns describing potential gene-expression changes across diverse tissue and cell types (**Figure 3F**). For detailed instructions and illustrative examples of each function, users can consult the Documentation page, which provides a step-by-step walkthrough with screenshots or short animation.

### A case study using ClinVar mutations

To demonstrate the performance of SomaMutDB annotations, we selected mutations from the ClinVar database (50), including one pathogenic group and one benign group. **Figure 4** (and **Figure S3**) shows the score distributions of these two groups for six representative annotation algorithms, plotted against the background distribution of all mutations in SomaMutDB. As expected, benign mutations exhibited substantially lower normalized scores than pathogenic mutations. Moreover, in most algorithms, pathogenic mutations were consistently shifted toward the extreme “pathogenic” end of the distribution relative to the SomaMutDB background, demonstrating the ability of the integrated framework to effectively distinguish between benign and pathogenic variants.

**Figure 4.**
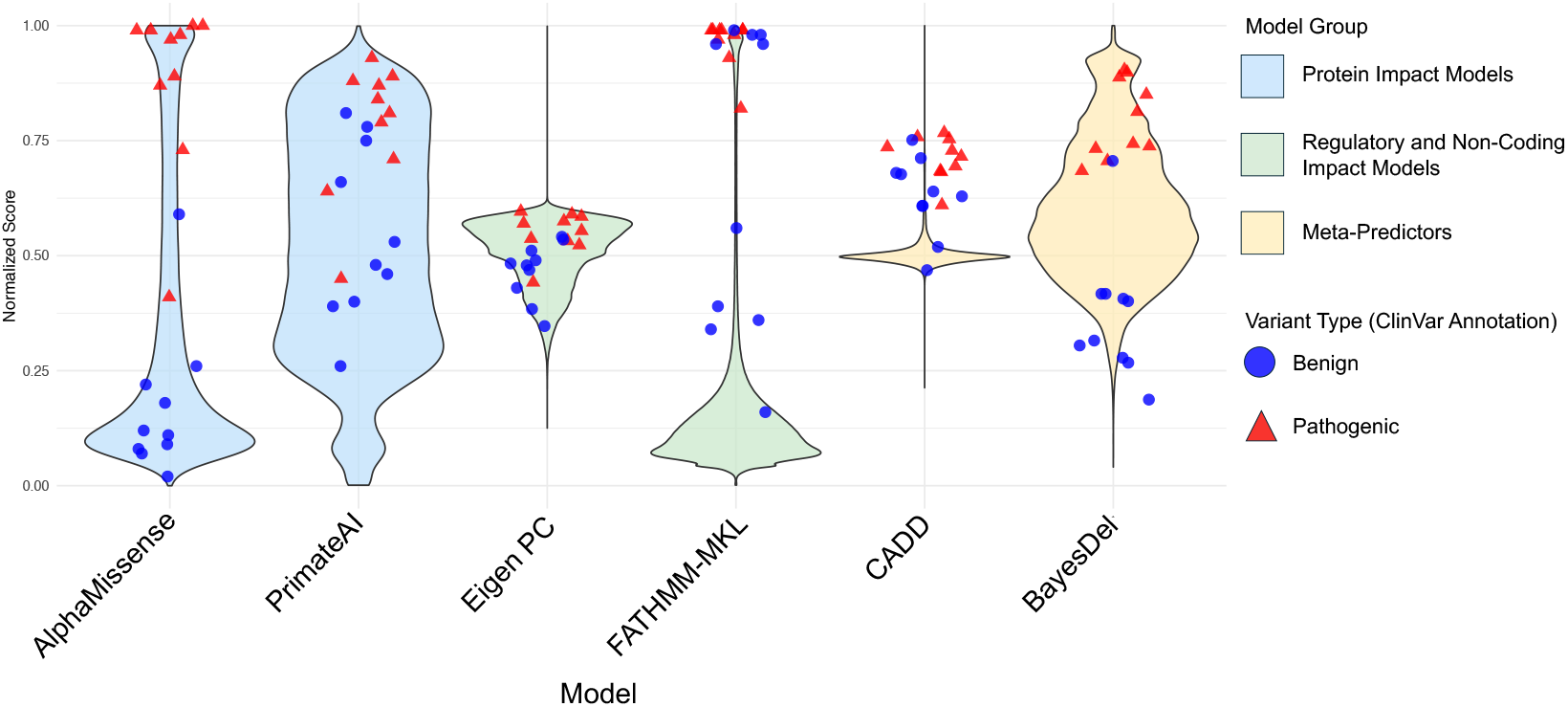
Benchmarking model predictions using ClinVar variants. Randomly selected ClinVar variants classified as benign or pathogenic (n=10 per group) were scored using six predictive models: two Protein/Sequence Impact Models (AlphaMissense and PrimateAI), two Regulatory/Non-Coding Impact Models (Eigen PC and FATHMM-MKL Non-Coding), and two Meta-Predictors (CADD and BayesDel-AddAF). Resulting ClinVar mutation scores were overlaid on the score distributions obtained from the full SomaMutDB database and labeled according to ClinVar annotation as either benign or pathogenic. Each plot illustrates the position of ClinVar variants relative to the overall distribution for the corresponding model.

## Conclusions and Perspectives

SomaMutDB 2.0 represents a major advance in the systematic study of somatic mutations in normal human tissues by combining large-scale data integration with a comprehensive functional annotation framework. With nearly nine million mutations curated across diverse tissues, age groups, and sequencing platforms, and annotated using 22 complementary predictive models, the database provides both breadth and depth for mutation discovery and interpretation. By enabling users not only to explore pre-annotated variants but also to upload and analyze their own mutations in the context of mutations from normal, non-diseased tissues, SomaMutDB 2.0 offers an improved framework for understanding mutational impact. Looking ahead, we plan to incorporate more disease- and tissue-specific annotation tools, such as DS-MVP (51), to further refine prediction accuracy. To enhance accessibility, we also envision developing an AI-powered assistant, such as DrBioRight 2.0 (52), to guide users in analyzing both database and user-supplied somatic mutation data. Ultimately, SomaMutDB aims to serve as a central and evolving platform for advancing our understanding of the landscape and biological impact of somatic mosaicism in human health and aging.

J.V. is a co-founder of Mutagentech Inc. Others declare no conflict of interest.

## Data Availability

All data from SomaMutDB 2.0 is available for download at: https://somamutdb.org/SomaMutDB/.

## Author Contributions Statement

Conceptualization: L.Z. & X.D.; Data curation: S.S.; Software: A.S., & S.S.; Validation: J.K.; Supervision: L.Z., J.V., & X.D.; Writing: All authors.

## References

1. Hutter, C. and Zenklusen, J.C. (2018) The Cancer Genome Atlas: Creating Lasting Value beyond Its Data. Cell, 173, 283–285.

2. Consortium, I.T.P.-C.A.o.W.G. (2020) Pan-cancer analysis of whole genomes. Nature, 578, 82–93.

3. Tate, J.G., Bamford, S., Jubb, H.C., Sondka, Z., Beare, D.M., Bindal, N., Boutselakis, H., Cole, C.G., Creatore, C., Dawson, E. et al. (2019) COSMIC: the Catalogue Of Somatic Mutations In Cancer. Nucleic Acids Res, 47, D941–d947.

4. Chen, C., Xing, D., Tan, L., Li, H., Zhou, G., Huang, L. and Xie, X.S. (2017) Single-cell whole-genome analyses by Linear Amplification via Transposon Insertion (LIANTI). Science, 356, 189–194.

5. Zhang, L., Lee, M., Maslov, A.Y., Montagna, C., Vijg, J. and Dong, X. (2024) Analyzing somatic mutations by single-cell whole-genome sequencing. Nat Protoc, 19, 487–516.

6. Cagan, A., Baez-Ortega, A., Brzozowska, N., Abascal, F., Coorens, T.H.H., Sanders, M.A., Lawson, A.R.J., Harvey, L.M.R., Bhosle, S., Jones, D. et al. (2022) Somatic mutation rates scale with lifespan across mammals. Nature.

7. Kapadia, C.D., Williams, N., Dawson, K.J., Watson, C., Yousefzadeh, M.J., Le, D., Nyamondo, K., Kodavali, S., Cagan, A., Waldvogel, S. et al. (2025) Clonal dynamics and somatic evolution of haematopoiesis in mouse. Nature, 641, 681–689.

8. Schmitt, M.W., Kennedy, S.R., Salk, J.J., Fox, E.J., Hiatt, J.B. and Loeb, L.A. (2012) Detection of ultra-rare mutations by next-generation sequencing. Proc Natl Acad Sci U S A, 109, 14508–14513.

9. Abascal, F., Harvey, L.M.R., Mitchell, E., Lawson, A.R.J., Lensing, S.V., Ellis, P., Russell, A.J.C., Alcantara, R.E., Baez-Ortega, A., Wang, Y. et al. (2021) Somatic mutation landscapes at single-molecule resolution. Nature, 593, 405–410.

10. Ganz, J., Luquette, L.J., Bizzotto, S., Miller, M.B., Zhou, Z., Bohrson, C.L., Jin, H., Tran, A.V., Viswanadham, V.V., McDonough, G. et al. (2024) Contrasting somatic mutation patterns in aging human neurons and oligodendrocytes. Cell, 187, 1955–1970.e1923.

11. Zhang, L., Dong, X., Lee, M., Maslov, A.Y., Wang, T. and Vijg, J. (2019) Single-cell whole-genome sequencing reveals the functional landscape of somatic mutations in B lymphocytes across the human lifespan. Proc Natl Acad Sci U S A, 116, 9014–9019.

12. Vrtačnik, P., Merino, L.G., Subhash, S., Helgadóttir, H.T., Bardin, M., Stefani, F., Wang, D., Chen, P., Franco, I., Revêchon, G. et al. (2025) Induced somatic mutation accumulation during skeletal muscle regeneration reduces muscle strength. Nat Aging.

13. Sun, S., Wang, Y., Maslov, A.Y., Dong, X. and Vijg, J. (2022) SomaMutDB: a database of somatic mutations in normal human tissues. Nucleic Acids Res, 50, D1100–d1108.

14. Huang, Z., Sun, S., Lee, M., Maslov, A.Y., Shi, M., Waldman, S., Marsh, A., Siddiqui, T., Dong, X., Peter, Y. et al. (2022) Single-cell analysis of somatic mutations in human bronchial epithelial cells in relation to aging and smoking. Nat Genet, 54, 492–498.

15. Shea, A., Bartz, J., Zhang, L. and Dong, X. (2023) Predicting mutational function using machine learning. Mutat Res Rev Mutat Res, 791, 108457.

16. Coorens, T.H.H., Oh, J.W., Choi, Y.A., Lim, N.S., Zhao, B., Voshall, A., Abyzov, A., Antonacci-Fulton, L., Aparicio, S., Ardlie, K.G. et al. (2025) The Somatic Mosaicism across Human Tissues Network. Nature, 643, 47–59.

17. Lawson, A.R.J., Abascal, F., Nicola, P.A., Lensing, S.V., Roberts, A.L., Kalantzis, G., Baez-Ortega, A., Brzozowska, N., El-Sayed Moustafa, J.S., Vaitkute, D. et al. (2024) Somatic mutation and selection at epidemiological scale. medRxiv, 2024.2010.2030.24316422.

18. Maslov, A.Y., Makhortov, S., Sun, S., Heid, J., Dong, X., Lee, M. and Vijg, J. (2022) Single-molecule, quantitative detection of low-abundance somatic mutations by high-throughput sequencing. Sci Adv, 8, eabm3259.

19. Vaser, R., Adusumalli, S., Leng, S.N., Sikic, M. and Ng, P.C. (2016) SIFT missense predictions for genomes. Nat Protoc, 11, 1–9.

20. Rentzsch, P., Witten, D., Cooper, G.M., Shendure, J. and Kircher, M. (2019) CADD: predicting the deleteriousness of variants throughout the human genome. Nucleic Acids Res, 47, D886–d894.

21. Cheng, J., Novati, G., Pan, J., Bycroft, C., Žemgulytė, A., Applebaum, T., Pritzel, A., Wong, L.H., Zielinski, M., Sargeant, T. et al. (2023) Accurate proteome-wide missense variant effect prediction with AlphaMissense. Science, 381, eadg7492.

22. Frazer, J., Notin, P., Dias, M., Gomez, A., Min, J.K., Brock, K., Gal, Y. and Marks, D.S. (2021) Disease variant prediction with deep generative models of evolutionary data. Nature, 599, 91–95.

23. Sundaram, L., Gao, H., Padigepati, S.R., McRae, J.F., Li, Y., Kosmicki, J.A., Fritzilas, N., Hakenberg, J., Dutta, A., Shon, J. et al. (2018) Predicting the clinical impact of human mutation with deep neural networks. Nat Genet, 50, 1161–1170.

24. Miller, M.B., Huang, A.Y., Kim, J., Zhou, Z., Kirkham, S.L., Maury, E.A., Ziegenfuss, J.S., Reed, H.C., Neil, J.E., Rento, L. et al. (2022) Somatic genomic changes in single Alzheimer’s disease neurons. Nature, 604, 714–722.

25. Nguyen, L., Jager, M., Lieshout, R., de Ruiter, P.E., Locati, M.D., Besselink, N., van der Roest, B., Janssen, R., Boymans, S., de Jonge, J. et al. (2021) Precancerous liver diseases do not cause increased mutagenesis in liver stem cells. Commun Biol, 4, 1301.

26. de Kanter, J.K., Peci, F., Bertrums, E., Rosendahl Huber, A., van Leeuwen, A., van Roosmalen, M.J., Manders, F., Verheul, M., Oka, R., Brandsma, A.M. et al. (2021) Antiviral treatment causes a unique mutational signature in cancers of transplantation recipients. Cell Stem Cell, 28, 1726–1739.e1726.

27. Kuijk, E., Kranenburg, O., Cuppen, E. and Van Hoeck, A. (2022) Common anti-cancer therapies induce somatic mutations in stem cells of healthy tissue. Nat Commun, 13, 5915.

28. Rouhani, F.J., Zou, X., Danecek, P., Badja, C., Amarante, T.D., Koh, G., Wu, Q., Memari, Y., Durbin, R., Martincorena, I. et al. (2022) Substantial somatic genomic variation and selection for BCOR mutations in human induced pluripotent stem cells. Nat Genet, 54, 1406–1416.

29. Olafsson, S., Rodriguez, E., Lawson, A.R.J., Abascal, F., Huber, A.R., Suembuel, M., Jones, P.H., Gerdes, S., Martincorena, I., Weidinger, S. et al. (2023) Effects of psoriasis and psoralen exposure on the somatic mutation landscape of the skin. Nat Genet, 55, 1892–1900.

30. Wang, H.T., Zhao, L., Yang, L.Q., Ge, M.X., Yang, X.L., Gao, Z.L., Cun, Y.P., Xiao, F.H. and Kong, Q.P. (2024) Scrutiny of genome-wide somatic mutation profiles in centenarians identifies the key genomic regions for human longevity. Aging Cell, 23, e13916.

31. Zhou, J., Theesfeld, C.L., Yao, K., Chen, K.M., Wong, A.K. and Troyanskaya, O.G. (2018) Deep learning sequence-based ab initio prediction of variant effects on expression and disease risk. Nat Genet, 50, 1171–1179.

32. McLaren, W., Gil, L., Hunt, S.E., Riat, H.S., Ritchie, G.R., Thormann, A., Flicek, P. and Cunningham, F. (2016) The Ensembl Variant Effect Predictor. Genome Biol, 17, 122.

33. Liu, X., Li, C., Mou, C., Dong, Y. and Tu, Y. (2020) dbNSFP v4: a comprehensive database of transcript-specific functional predictions and annotations for human nonsynonymous and splice-site SNVs. Genome Med, 12, 103.

34. Quang, D., Chen, Y. and Xie, X. (2015) DANN: a deep learning approach for annotating the pathogenicity of genetic variants. Bioinformatics, 31, 761–763.

35. Raimondi, D., Tanyalcin, I., Ferté, J., Gazzo, A., Orlando, G., Lenaerts, T., Rooman, M. and Vranken, W. (2017) DEOGEN2: prediction and interactive visualization of single amino acid variant deleteriousness in human proteins. Nucleic Acids Res, 45, W201–w206.

36. Samocha, K.E., Kosmicki, J.A., Karczewski, K.J., O’Donnell-Luria, A.H., Pierce-Hoffman, E., MacArthur, D.G., Neale, B.M. and Daly, M.J. (2017) Regional missense constraint improves variant deleteriousness prediction. bioRxiv, 148353.

37. Ionita-Laza, I., McCallum, K., Xu, B. and Buxbaum, J.D. (2016) A spectral approach integrating functional genomic annotations for coding and noncoding variants. Nat Genet, 48, 214–220.

38. Lu, Q., Hu, Y., Sun, J., Cheng, Y., Cheung, K.H. and Zhao, H. (2015) A statistical framework to predict functional non-coding regions in the human genome through integrated analysis of annotation data. Sci Rep, 5, 10576.

39. Malhis, N., Jacobson, M., Jones, S.J.M. and Gsponer, J. (2020) LIST-S2: taxonomy based sorting of deleterious missense mutations across species. Nucleic Acids Res, 48, W154–w161.

40. Feng, B.J. (2017) PERCH: A Unified Framework for Disease Gene Prioritization. Hum Mutat, 38, 243–251.

41. Jagadeesh, K.A., Wenger, A.M., Berger, M.J., Guturu, H., Stenson, P.D., Cooper, D.N., Bernstein, J.A. and Bejerano, G. (2016) M-CAP eliminates a majority of variants of uncertain significance in clinical exomes at high sensitivity. Nat Genet, 48, 1581–1586.

42. Li, C., Zhi, D., Wang, K. and Liu, X. (2022) MetaRNN: differentiating rare pathogenic and rare benign missense SNVs and InDels using deep learning. Genome Med, 14, 115.

43. Dong, C., Wei, P., Jian, X., Gibbs, R., Boerwinkle, E., Wang, K. and Liu, X. (2015) Comparison and integration of deleteriousness prediction methods for nonsynonymous SNVs in whole exome sequencing studies. Hum Mol Genet, 24, 2125–2137.

44. Qi, H., Zhang, H., Zhao, Y., Chen, C., Long, J.J., Chung, W.K., Guan, Y. and Shen, Y. (2021) MVP predicts the pathogenicity of missense variants by deep learning. Nat Commun, 12, 510.

45. Ioannidis, N.M., Rothstein, J.H., Pejaver, V., Middha, S., McDonnell, S.K., Baheti, S., Musolf, A., Li, Q., Holzinger, E., Karyadi, D. et al. (2016) REVEL: An Ensemble Method for Predicting the Pathogenicity of Rare Missense Variants. Am J Hum Genet, 99, 877–885.

46. Shihab, H.A., Rogers, M.F., Gough, J., Mort, M., Cooper, D.N., Day, I.N., Gaunt, T.R. and Campbell, C. (2015) An integrative approach to predicting the functional effects of non-coding and coding sequence variation. Bioinformatics, 31, 1536–1543.

47. Mao, Y., Chen, H., Liang, H., Meric-Bernstam, F., Mills, G.B. and Chen, K. (2013) CanDrA: cancer-specific driver missense mutation annotation with optimized features. PLoS One, 8, e77945.

48. Moyon, L., Berthelot, C., Louis, A., Nguyen, N.T.T. and Roest Crollius, H. (2022) Classification of non-coding variants with high pathogenic impact. PLoS Genet, 18, e1010191.

49. Steinhaus, R., Proft, S., Schuelke, M., Cooper, D.N., Schwarz, J.M. and Seelow, D. (2021) MutationTaster2021. Nucleic Acids Res, 49, W446–w451.

50. Landrum, M.J., Chitipiralla, S., Kaur, K., Brown, G., Chen, C., Hart, J., Hoffman, D., Jang, W., Liu, C., Maddipatla, Z. et al. (2025) ClinVar: updates to support classifications of both germline and somatic variants. Nucleic Acids Res, 53, D1313–d1321.

51. Chen, Q., Quan, L., Cao, L., Zhang, B., Zhang, Z., Peng, L., Wang, J., Jiang, Y., Nie, L., Li, G. et al. (2025) DS-MVP: identifying disease-specific pathogenicity of missense variants by pre-training representation. Brief Bioinform, 26.

52. Liu, W., Li, J., Tang, Y., Zhao, Y., Liu, C., Song, M., Ju, Z., Kumar, S.V., Lu, Y., Akbani, R. et al. (2025) DrBioRight 2.0: an LLM-powered bioinformatics chatbot for large-scale cancer functional proteomics analysis. Nat Commun, 16, 2256.

