## Supplementary Figures for "SomaMutDB 2.0: A comprehensive database for exploring somatic mutations and their functional impact in normal human tissues"

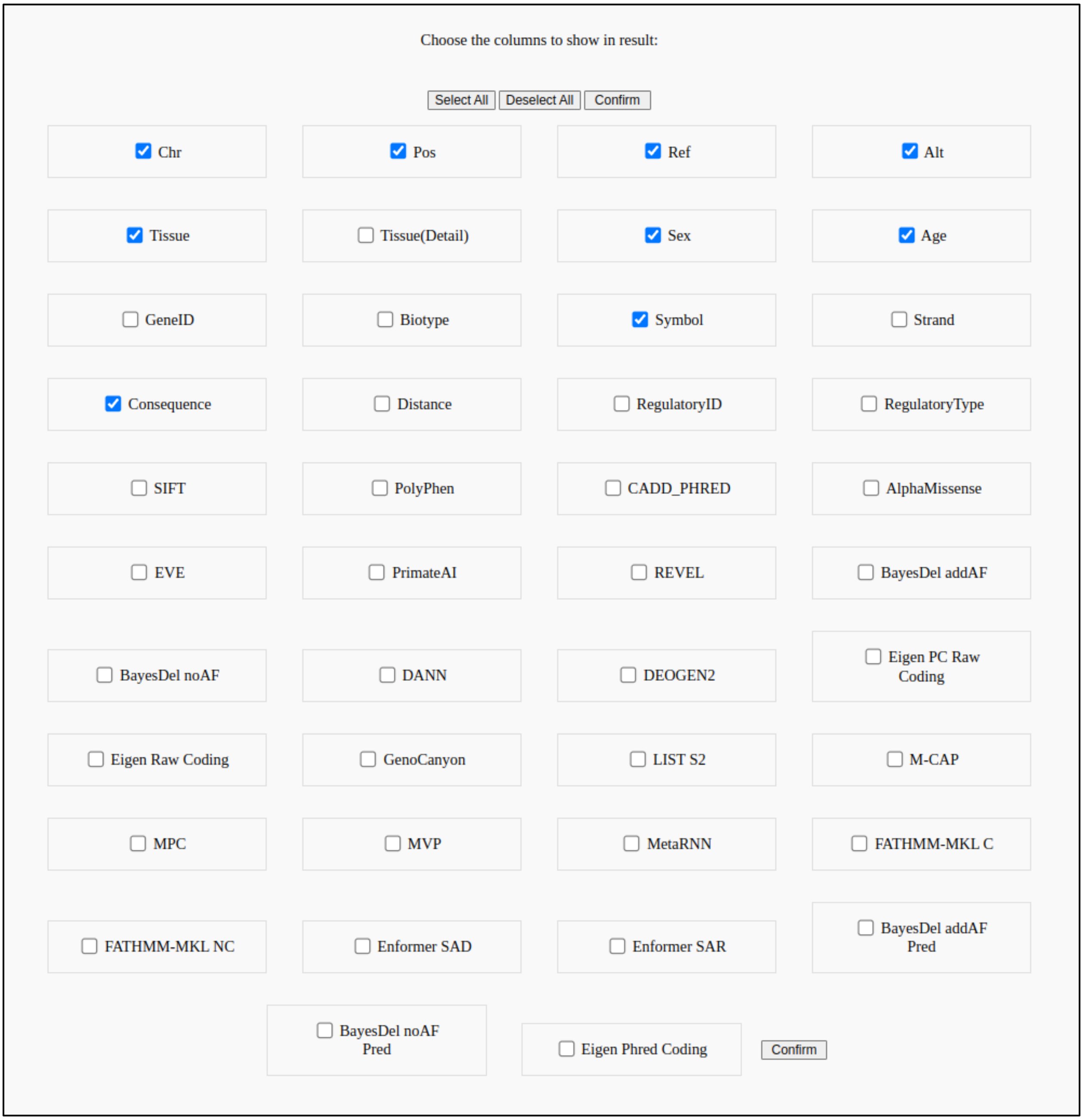


**Figure S1. Screenshot of updated column selection for search feature.**

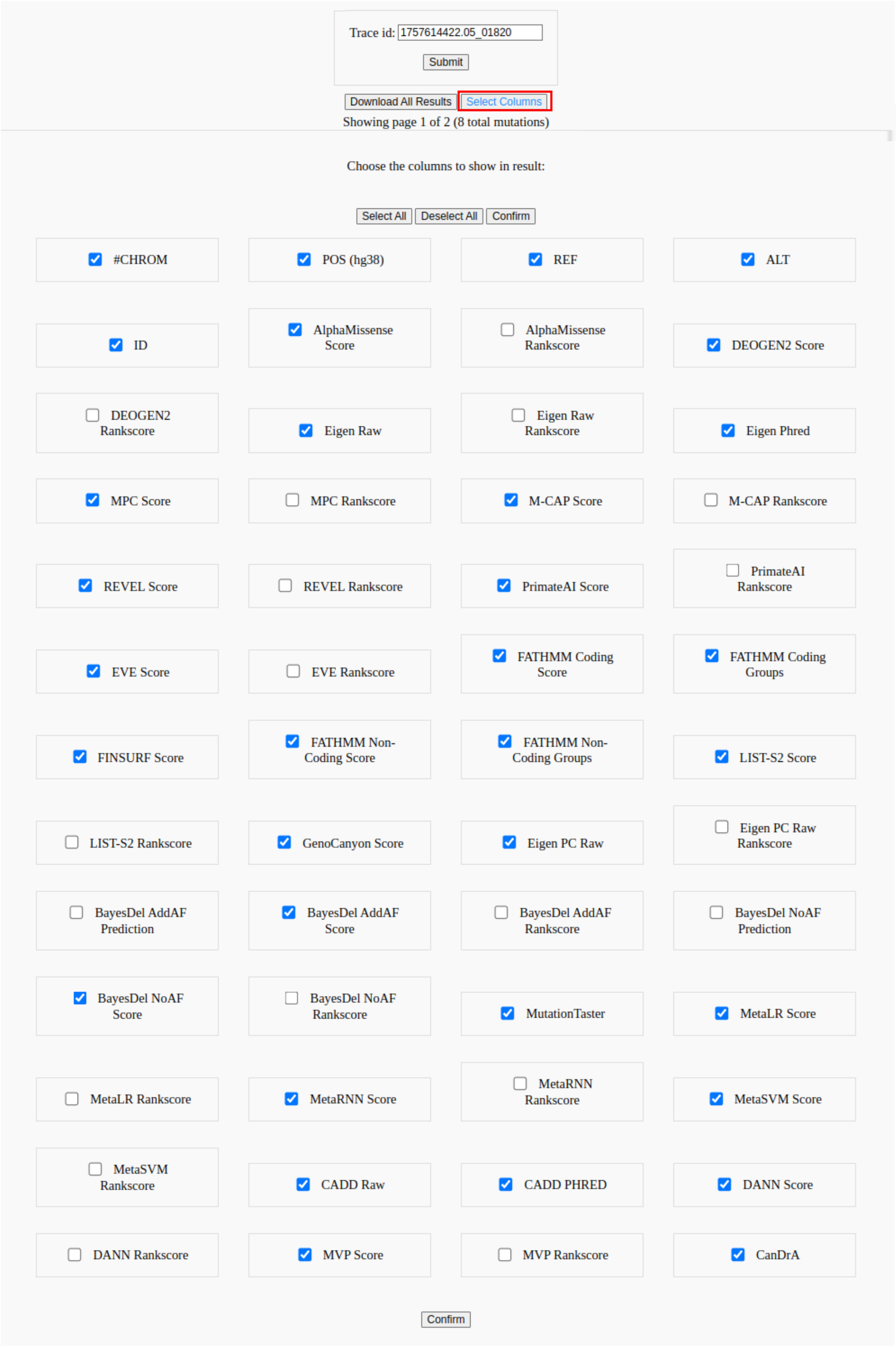


**Figure S2. Screenshot of updated column selection for pathogenic prediction results.**

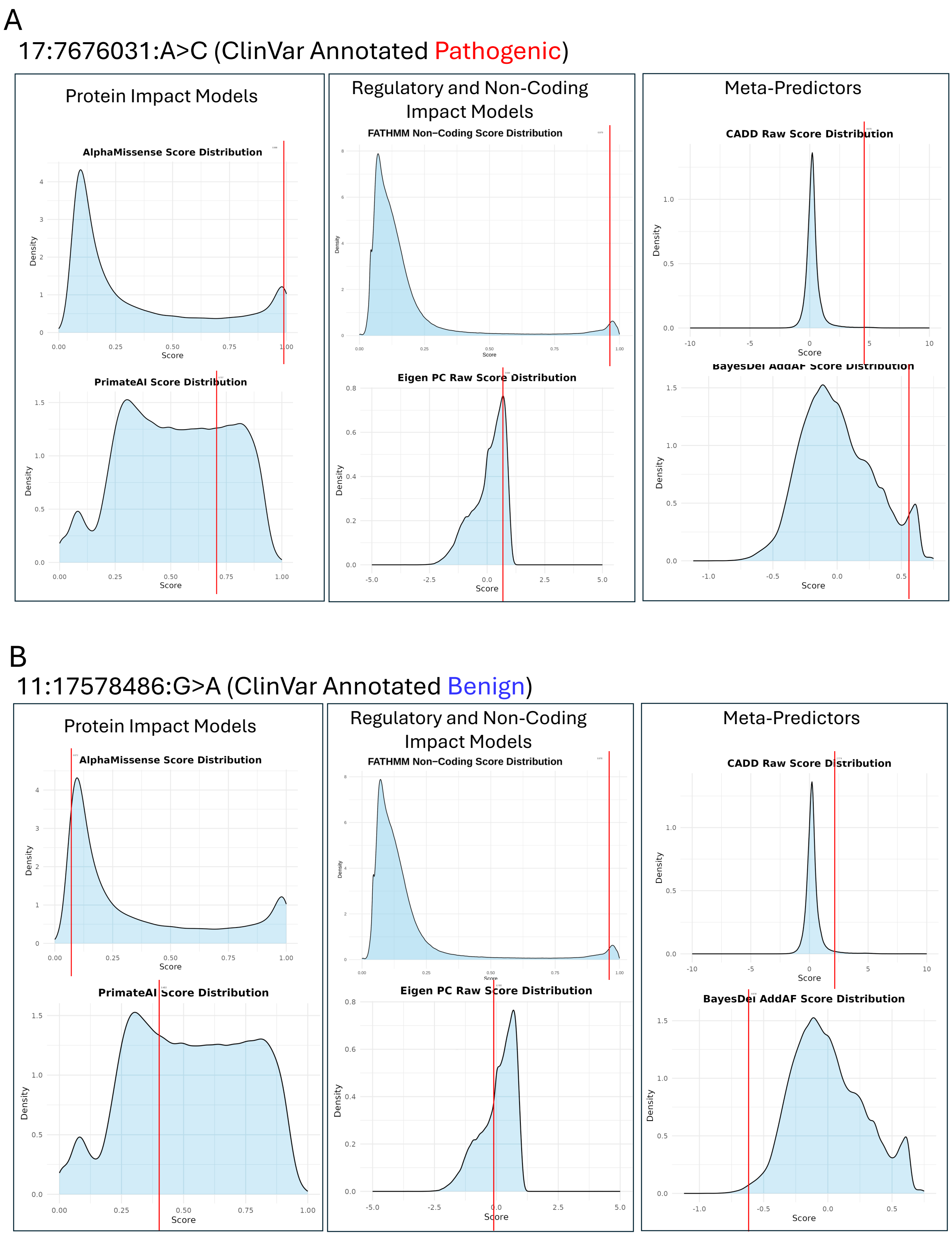


**Figure S3. Density plot view as displayed in the SomaMutDB website.**
